# A Novel Mouse Model of Chronic Primary Pain Conditions that Integrates Clinically Relevant Genetic and Environmental Factors

**DOI:** 10.1101/2022.12.18.520949

**Authors:** Yaomin Wang, Shin Hyung Kim, Marguerita E Klein, Jiegen Chen, Elizabeth Gu, Shad B. Smith, Andrey Bortsov, Gary D Slade, Xin Zhang, Andrea G Nackley

## Abstract

Chronic primary pain conditions (CPPCs) affect over 100 million people, predominantly women. Yet, they remain ineffectively treated due, in large part, to lack of valid animal models with translational relevance. Here, we characterized a novel mouse model of CPPCs that integrated clinically-relevant genetic (catechol-o-methyltransferase; COMT knockdown) and environmental (stress and minor injury) factors. Compared to wildtype mice, COMT+/- mice undergoing the repeated swim stress and molar extraction surgery intervention exhibited pronounced multi-site body pain and depressive-like behavior lasting more than 3 months. The COMT+/- mice undergoing the intervention also exhibited enhanced activity of primary afferent DRG nociceptors innervating hindpaw and back sites and increased plasma levels of norepinephrine and the pro-inflammatory cytokines IL-6 and IL-17A. Notably, the pain and depressive-like behavior was of greater magnitude and longer duration (lasting at least 12 months) in females compared to males. Further, increases in anxiety-like behavior and IL-6 levels were female-specific. Intervention-induced body pain and nociception in COMT+/- mice was blocked by a beta-3 adrenergic antagonist, demonstrating predictive validity. Finally, the effect of COMT genotype x stress interactions on pain and IL-6 and IL-17A levels was observed in our clinical CPPC case-control cohort, demonstrating construct validity. Thus, our novel mouse model reliably recapitulates clinically- and biologically-relevant features of CPPCs and can be further implemented to test underlying mechanisms and discover new therapeutics.

**One Sentence Summary:** We developed a novel mouse model of chronic primary pain conditions that shares similar genetic, biologic, and clinical features of patients.

## INTRODUCTION

Chronic primary pain conditions (CPPC_S_) including fibromyalgia syndrome (FMS), temporomandibular disorder (TMD), tension-type headache (TTH), irritable bowel syndrome (IBS), vestibulodynia (VBD), and low back pain (LBP) affect nearly 1 in every 3 Americans, predominantly females (*1, 2*). These conditions are characterized by persistent pain in the absence of tissue damage and often co-occur, thereby affecting multiple body sites. In fact, a recent community-based study of 655 adults in the United States found that overlap is the ‘norm’ for these conditions and, while the magnitude of overlap varies for each, the range may be as high as 50 to 90% (*3*). Overlapping pain creates serious problems for patients, adding to the suffering and disability caused by a single pain condition (*4*). Healthcare providers who manage one type of chronic pain often are at a loss to manage pain occurring elsewhere in the body. Despite their high prevalence and significant economic burden, CPPCs remain ineffectively treated due, in large part, to the lack of valid animal models with translational relevance to explore underlying mechanisms and screen for effective therapies.

CPPCs have been challenging to model due to their idiopathic pathogenesis (*5*). While the etiology of CPPCs is complex, accumulating evidence suggests that the origin of the pain is linked to interactions in genetic and environmental events resulting in heightened catecholamine bioavailability and systemic inflammation. Patients with CPPC_S_ have increased levels of catecholamines in circulation (*6-9*) alongside reduced levels of catechol-O-methyltransferase (COMT) (*10, 11*), a ubiquitously expressed enzyme that metabolizes catecholamines (*11, 12*). An estimated 66% of patients with CPPCs have functional variants in the *COMT* gene that result in low COMT activity (*11, 13, 14*). The ‘low COMT activity’ variants are associated with increased CPPC onset (*15-19*), increased pain in response to experimental stimuli (*20*), and increased pain-related comorbidities such as depression and anxiety (*21-23*). For example, individuals with the low COMT activity genotype report enhanced pain following stressful events (eg, psychological strain) (*24, 25*) and injurious surgical procedures (eg, molar extraction) (*26-29*). Low COMT (*30*), stress (*31*), and injury (*32*) can produce pain by increasing the production of pro-inflammatory cytokines that sensitize nociceptors (*33-36*). Pro-inflammatory cytokine levels are elevated in CPPCs such as TMD (*37-41*), TTH (*42-44*), IBS (*45-47*), VBD (*46, 48, 49*), and FMS (*50-54*). Further, they are correlated with pain intensity (*39, 41, 51, 55, 56*), perceived stress (*57*), and mood disorders (*58, 59*) that often accompany CPPCs. Collectively, these findings from clinical studies point to critical factors that should be considered in developing improved preclinical animal models, including 1) low COMT activity genotype, environmental stressors, and minor injuries as independent variables, 2) multi-site body pain and comorbid depression/anxiety as dependent variables, and 3) circulating cytokines as pain-relevant biomarkers.

Herein, we established a genetic x environmental approach in which *COMT+/-* mice are exposed to a 3-day swim stress paradigm followed by a minor molar extraction surgery. Compared to wildtype mice, we found that *COMT+/-* mice, which exhibit normal baseline behavioral responses, develop exaggerated multi-site body pain and comorbid depressive behavior which is of greater magnitude and longer duration in females compared to males. Further, *COMT+/-* mice exposed to the intervention exhibit increased activity of primary afferent nociceptors and increased plasma levels of the pro-inflammatory cytokines IL-6 and IL-17A. Finally, we validated the effects of the COMT by stress interaction on pain and IL-6 and IL-17A cytokine levels in a human CPPC case control cohort.

## RESULTS

### COMT+/- mice undergoing swim stress and molar extraction surgery exhibit robust mechanical pain at multiple body sites

To model CPPCs, we employed COMT+/- mice that carry a genetic predisposition towards pain, but otherwise appear similar to wildtype mice, and exposed them to a 3-day repeated swim stress paradigm followed by a molar extraction surgery. The repeated swim stress model was used because it produces behavioral and physical arousal, hyperalgesia lasting around 7 days (*60*), sensitization of spinal sensory neurons (*61*), and the associated pain is mitigated by pharmacotherapies for stress-related disorders (*62*). Molar extraction surgery was used because it has been shown to increase risk for TMD (*26, 27*), especially among individuals with the low COMT activity genetic variants (*63*). We found that *COMT+/-* mice exposed to swim stress alone exhibited moderate mechanical hyperalgesia at plantar sites that resolved by day 7, while those undergoing molar extraction alone did not exhibit any change from baseline. Meanwhile, *COMT+/-* mice exposed to both swim stress and molar extraction exhibited severe mechanical hyperalgesia at plantar sites lasting at least 28 days (**Fig. S1**). After establishing the independent and joint effects of stress and injury on plantar hypersensitivity in COMT+/- mice, we proceeded to characterize multi-site body pain in COMT+/- compared to WT mice. Separate groups of mice were exposed to the swim stress and molar extraction surgery intervention or sham procedures and mechanical pain sensitivity measured at plantar, abdominal, and back sites over 28 days (**Fig. 1A**). Compared to WT Sham mice, the female and male WT mice exposed to the intervention exhibited mechanical allodynia **(Fig. 1B,D)** and hyperalgesia **(Fig. 1C,E)** at plantar sites lasting 28 days. Notably, the magnitude of the pain in both female and male *COMT+/-* mice exposed to the intervention was significantly greater than that of the WT mice exposed to the intervention, consistent with findings from clinical studies.

**Fig. 1.**
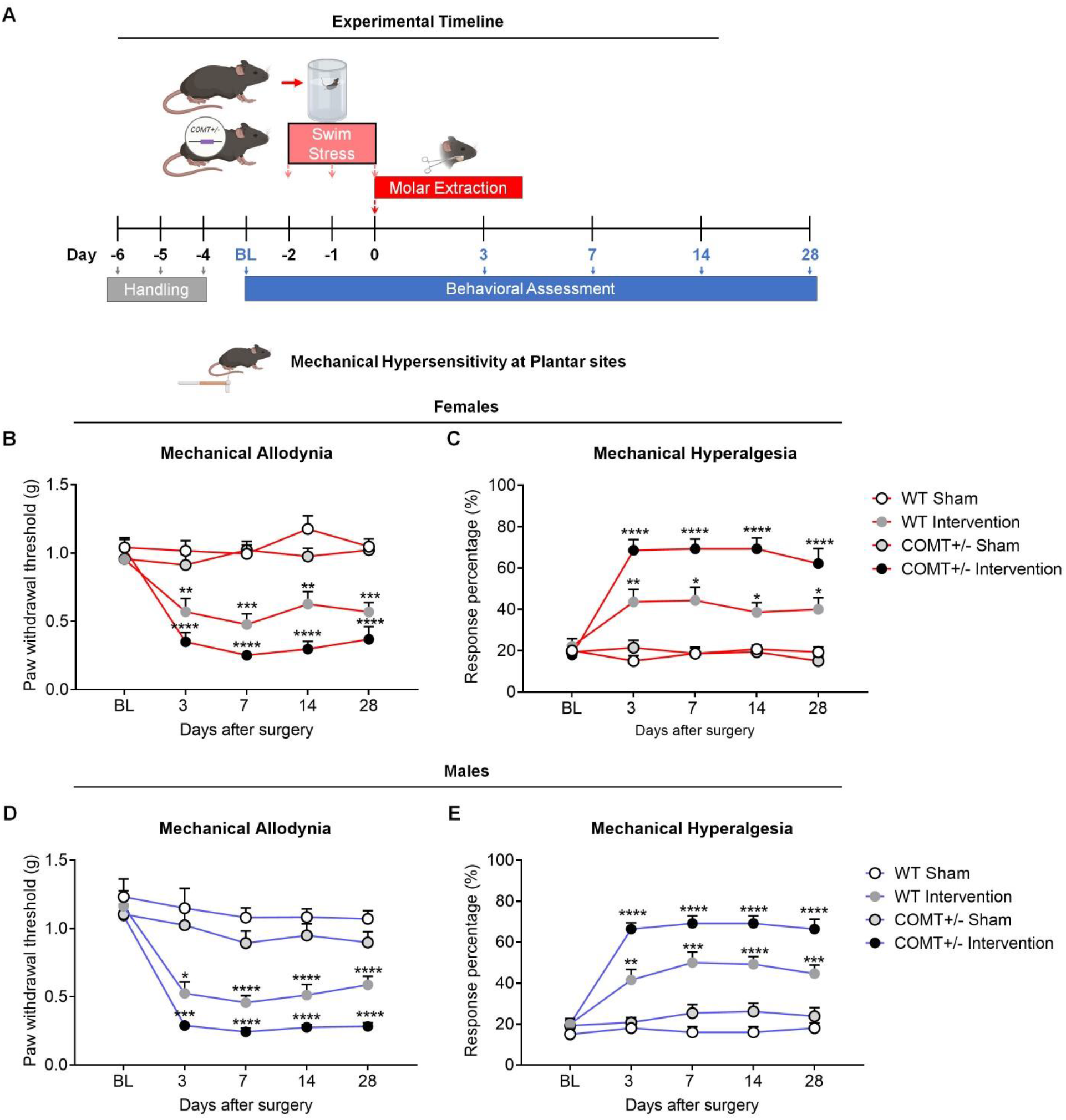
*COMT+/-* mice exposed to transient stress and injury exhibit robust and sustained mechanical hypersensitivity at plantar sites. **(A)** Experimental timeline for establishing the CPPC model and testing pain behaviors. **(B-C)** Both WT and *COMT+/-* female mice undergoing the intervention exhibit mechanical allodynia and hyperalgesia compared to sham controls, but the magnitude of the pain is greater in *COMT+/-* mice. **(D-E)** Similarly, both WT and COMT+/- male mice undergoing the intervention exhibit mechanical allodynia and hyperalgesia compared to sham controls, but the magnitude of the pain is greater in COMT+/- mice. *N* = 13-14 F and 10-15 M mice per group. Data represent mean ± SEM. ^*^*P <* 0.05, ^**^*P <* 0.01, ^***^*P* < 0.001, ^****^*P* < 0.0001 versus Sham Intervention. ^#^*P <* 0.05, ^##^*P* < 0.01, ^###^*P* < 0.001 versus WT Intervention.

Next, we tested mechanical sensitivity at remote sites using a spinal needle applied to the abdomen or back. The spinal needle tip was applied to each shaved, marked site 15 times and the number and type (normal/innocuous, noxious, or null) of responses recorded. When testing abdominal sites, we found that female *COMT+/-* intervention mice exhibited robust increases in the number of noxious withdrawals and decreases in null responses over 28 days compared to the other groups (**Fig. 2A-C**). In contrast, the male *COMT+/-* intervention mice only exhibited transient increases in the number of noxious withdrawals on day 7 (**Fig. 2D-F**). Similarly, when testing back sites, we found that female COMT+/- intervention mice exhibited robust increases in the number of noxious responses and decreases in null responses at back sites over 28 days (**Fig. 2 G-I**), while male COMT+/- intervention mice exhibited transient increases in the number of noxious withdrawls on days 3-14 (**Fig. 2J-L**). Of note, both female and male mice in the WT intervention group exhibited consistent, though moderate, increases in noxious responses at abdominal and back sites on day 7. These data show that COMT+/- mice exposed to the stress and surgery intervention have robust mechanical pain at multiple body sites and, consistent with clinical syndromes, the pain is of greater magnitude and longer duration in females.

**Fig. 2.**
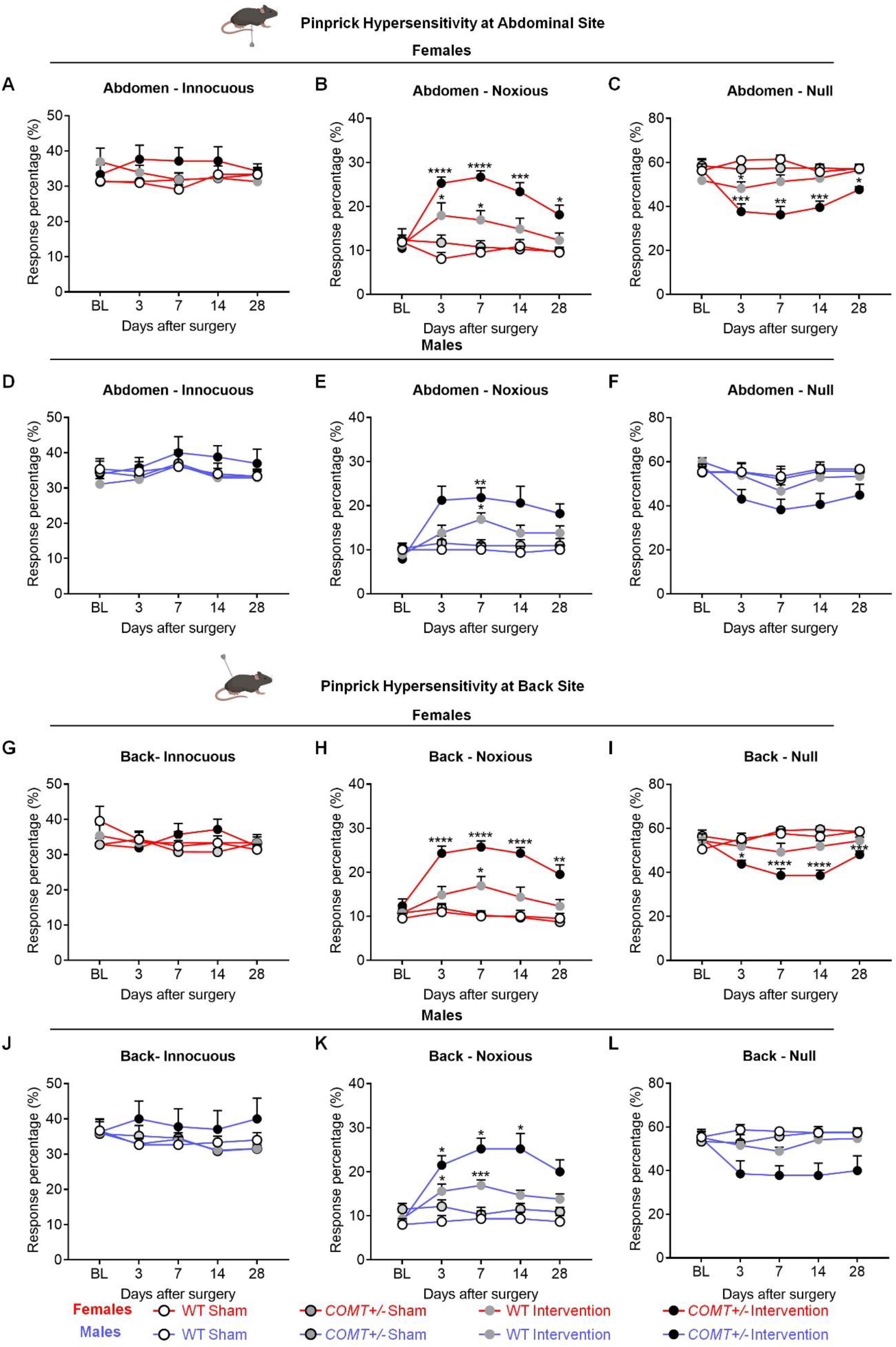
*COMT+/-* mice exposed to transient stress and injury exhibit robust and sustained mechanical hypersensitivity at abdominal and back sites. Compared to WT and sham control mice, female and male COMT+/- mice undergoing the intervention have greater mechanical hyperalgesia at **(A-F)** abdominal, and **(G-L)** back sites. Notably, females exhibit a greater magnitude of pinprick hyperalgesia compared to males. *N* = 13-14 F and 10-15 M mice per group. Data represent mean ± SEM. ^*^*P <* 0.05, ^**^*P <* 0.01, ^***^*P* < 0.001, ^****^*P* < 0.0001 versus Sham Intervention. ^#^*P <* 0.05, ^##^*P* < 0.01 WT intervention versus *COMT+/-* intervention.

### COMT+/- mice undergoing swim stress and molar extraction surgery exhibit peripheral sensitization

WT or COMT+/- mice were crossed with Pirt-GCaMP3 mice that express a calcium indicator exclusively in peripheral nociceptors, in order to examine the activity of primary afferent DRG neurons innervating the paw and back using *in vivo* Ca^2+^ imaging (**Fig. 3 A,B**). Compared to their respective sham control groups, both Pirt-GCaMP3 (defined as WT intervention) and Pirt-GCaMP3:COMT+/- (defined as *COMT+/-* intervention) groups exhibited significant pinch-evoked increases in the number of L4 DRG neurons innervating the hindpaw on day 7, with the largest increase observed for the COMT+/- intervention group (**Fig. 3C**). When evaluating the responses of L5 neurons innervating the back, we found that only the COMT+/- intervention group exhibited significant spinal needle-evoked increases on day 7 compared to controls. Though, the WT intervention group exhibited a trend towards increased L5 neuronal responses (**Fig. 3D**). These data suggest that our novel CPPC mice have developed widespread peripheral sensitization, evidenced by increases in DRG nociceptor activity, coincident with increases in pain behavior.

**Fig. 3.**
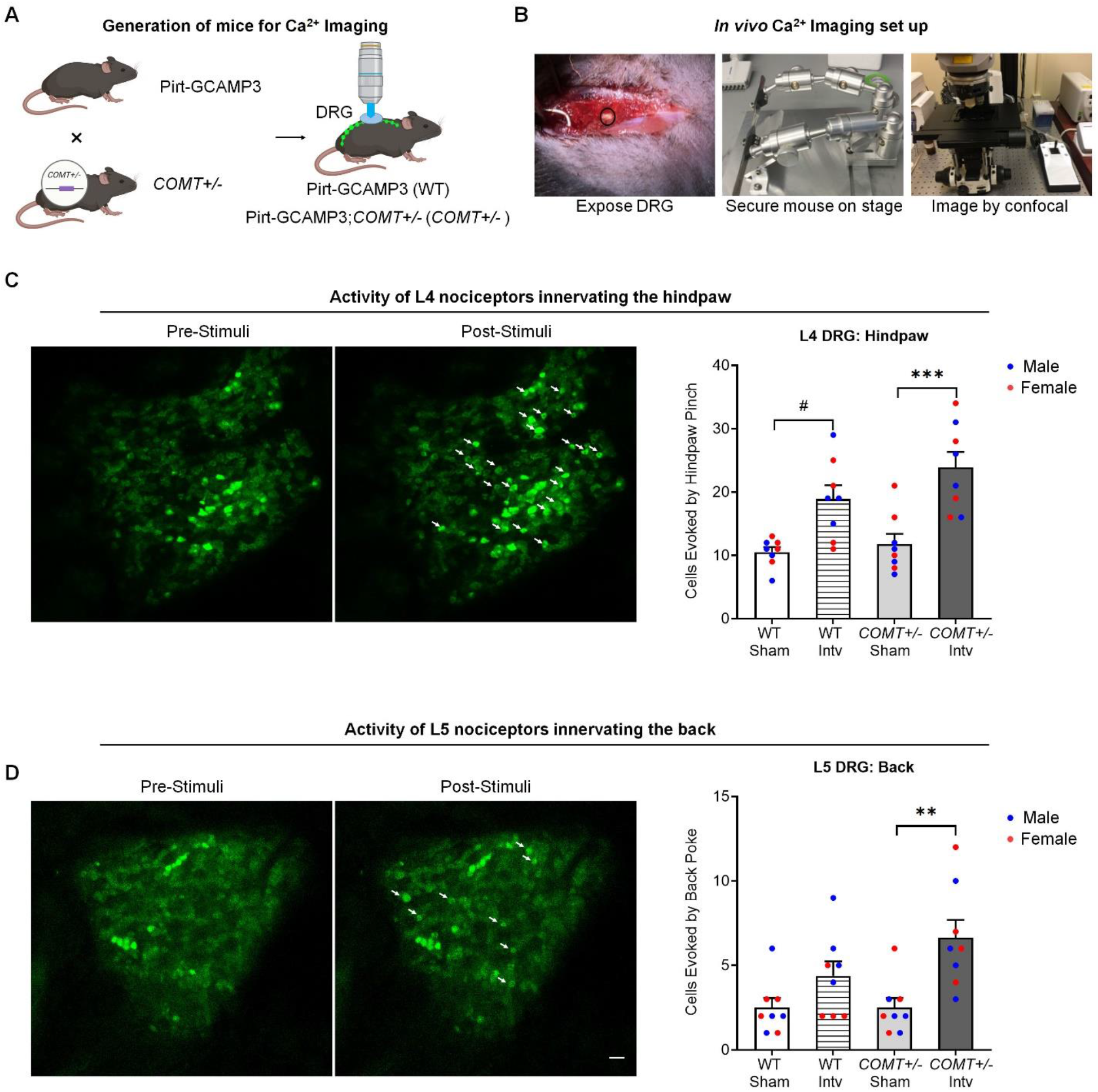
*COMT+/-* mice exposed to the intervention exhibit significant increases in the activity of nociceptors innervating plantar and back sites. Schematic illustration of the **(A)** strategy to knock in GCAMP3 in *COMT+/-* mice and **(B)** experimental paradigm and set up for *in vivo* Ca^2+^ imaging. **(C)** Representative confocal images for nociceptor activity before and after mechanical hindpaw pinch. Quantification of these data demonstrate transient stress- and injury-induced increased nociception activity in DRG neurons in *COMT+/-* mice. **(D)** Representative confocal images for nociceptor activity before and after back stimulation with spinal needles, Scale bar = 100 µm. Quantification of these data demonstrate transient stress- and injury-induced increased nociception activity in DRG neurons in COMT+/- mice. *N* = 8 mice (4F and 4M) per group. Data represent mean ± SEM. ^#^*P <* 0.05 versus WT intervention. ^**^*P <* 0.05, ^***^*P <* 0.05 versus *COMT+/-* Intervention.

### CPPC mice exhibit increased circulating levels of catecholamines and pro-inflammatory cytokines

Patients with CPPCs have increased levels of catecholamines in circulation alongside decreased levels of COMT. Using a standard ELISA, we measured norepinephrine (NE) levels in plasma samples collected from WT and *COMT+/-* sham and intervention mice at different time points. Basal NE levels were comparable between *COMT+/-* and WT mice (**Fig. 4A**). Following the intervention, the COMT+/- mice exhibited significant increases in plasma NE levels on days 7 and 28 compared to all other groups (**Fig. 4A**). These data provide further evidence of physiological similarity between our mouse model and patients with CPPCs.

**Fig. 4.**
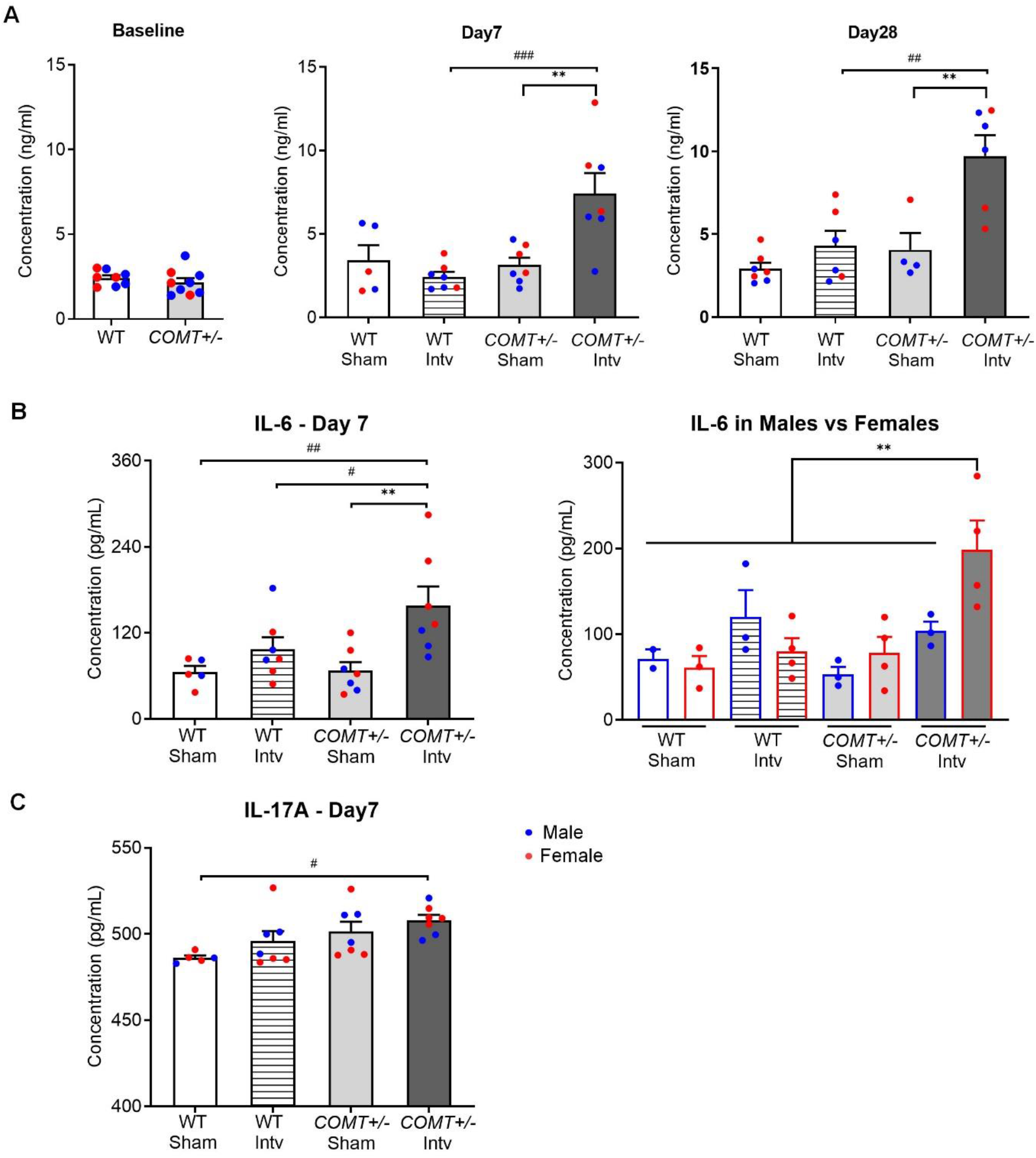
CPPC mice exhibit increased plasma levels of norepinephrine, IL-6, and IL-17. **(A)** Basal NE levels do not differ between WT and COMT+/- mice. Following the intervention, COMT+/- mice exhibit significant increases in NE on days 7 and 28. COMT+/- mice exposed to the intervention also exhibit increased levels of the pro-inflammatory cytokines **(B)** IL-6 and **(C)** IL-17A. Of note, IL-6 levels were increased in females, but not males **(D)**. *N* = 5–9 mice (3-4F and 2-6M) per group. Data represent mean ± SEM. ^**^*P <* 0.01, ^***^*P <* 0.001, versus Sham Intervention. ^#^*P* < 0.05, ^##^*P* < 0.01, ^###^*P* < 0.001 versus WT Intervention.

To identify putative biomarkers of CPPCs in our model, we used a Mesoscale Discovery multiplex platform to measure levels of 35 cytokines in plasma samples collected from the different experimental groups on day 7. Of these, IL-6 and IL-17A were significantly increased in the *COMT+/-* intervention group (**Fig. 4B,C**). Interestingly, the increase in IL-6 was specific to females, but not males (**Fig. 4D**). These data suggest that COMT genotype and stress/injury interactions regulate inflammation as well as pain.

### Mechanical pain sensitivity and nociceptor activity in CPPC mice is blocked by a β3AR antagonist

Beta-blockers that normalize catecholamine production have been shown to alleviate pain in patients with CPPCs (*64, 65*). In our prior work using a pharmacological model of COMT inhibition, we found that systematic delivery of the beta-3 adrenergic receptor (β3AR) antagonist SR59230A blocked OR486-induced mechanical hypersensitivity. To evaluate the analgesic efficacy of this β3AR antagonist in our novel CPPC model, separate groups of COMT+/- mice were implanted with osmotic minimpumps one day prior swim stress to deliver SR59230A (2 mg/kg/day or 10 mg/kg/day) or vehicle systematically over the course of 14 days (**Fig. 5A**). Results show that SR59230A attenuated the development of mechanical allodynia and mechanical hyperalgesia at plantar sites in a dose-dependent manner, with significant analgesia achieved by the higher 10 mg/kg/day dose (**Fig. 5B,C**). Importantly, this higher dose of SR59230A completely prevented the development of pain at abdominal and back sites, evidenced by the lack of change in noxious and null responses from baseline over the course of 28 days (**Fig. 5D,E**).

**Fig. 5.**
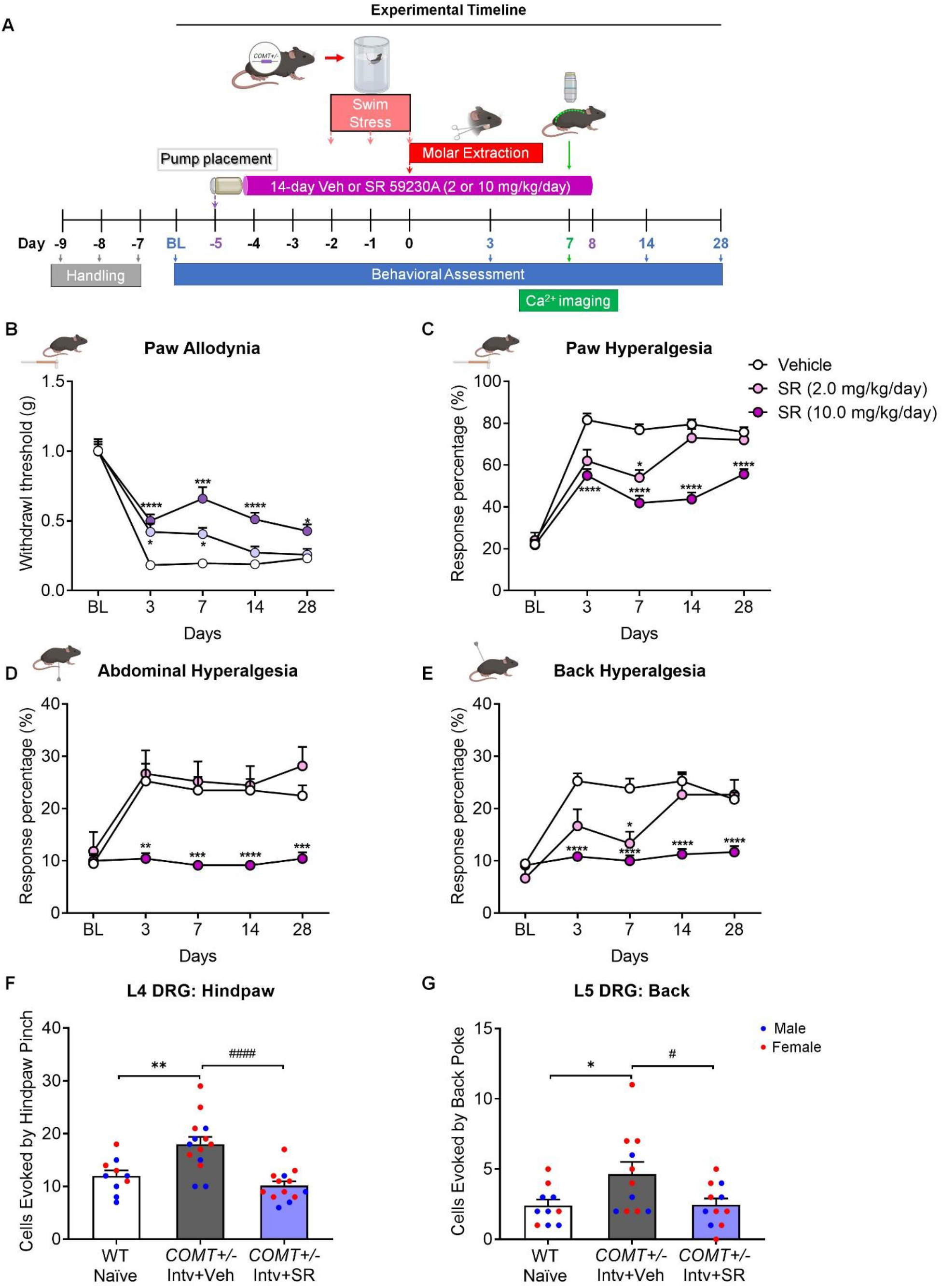
The β3AR antagonist SR59230A reduced multi-site mechanical hypersensitivity and nociceptor activity. **(A)** Experimental timeline for establishing the CPPC model and testing pain behaviors. Compared to naïve mice, SR59230A dose-dependently reduced mechanical hypersensitivity at **(B,C)** plantar, **(D)** abdominal, and **(E)** back sites among female and male CPPC mice. N = 19 (11M, 8F) in Vehicle, N=10 (5M, 5F) in SR (2.0 mg/kg/day), and N = 16 (7M, 9F) in SR (10.0 mg/kg/day) group. **(F,G)** SR59230A treatment prevented the activation of L4 and L5 DRG neurons in female and male CPPC mice. *N* = 10-14 mice per group. Data represent mean ± SEM. ^*^*P <* 0.05, ^**^*P <* 0.01, ^***^*P* < 0.001, ^****^*P* < 0.0001 versus vehicle or naïve. ^#^*P <* 0.05, ^*####*^*P <* 0.0001 versus *COMT+/-* Intervention+SR.

We further performed *in vivo* Ca^2+^ imaging on day 7 following the intervention to evaluate the effects of the β3AR antagonist on nociceptor activity. Results show that SR59230A completely blocked intervention-induced increases in the number of the L4 and L5 DRG neurons evoked by hind paw pinch (**Fig. 5F**) or back spinal needle poke (**Fig. 5G**).

### CPPC mice develop chronic pain and comorbid depression- and anxiety-like behaviors

To further characterize the time course of pain in our model, we employed a second cohort and tested them over the course of 48 weeks. Results show that both the female (**Fig. 6A-D**) and male (**Fig. 6E-H**) WT mice exposed to the intervention had ‘normal’ baseline level responses to mechanical stimuli at 2 months. Meanwhile, the *COMT+/-* intervention mice exhibited long lasting mechanical pain sensitivity. The female COMT+/- intervention mice demonstrated consistent and robust allodynia and hyperalgesia at plantar sites throughout the duration of the 48 week testing period. Interestingly the male COMT+/- mice exhibit pain at plantar sites for 28-32 weeks, after which time their pain progressively resolved. We also observed that abdominal and back pain lasts 8-12 weeks in female *COMT+/-* mice, but not the male COMT+/- mice. Thus, the pain in our model, which lasts at least 48 weeks (equivalent of 12 months) in female and 28 weeks in male mice (equivalent of 7 months) is in line with the IASP definition of chronic pain for humans (pain lasting ≥3 months) and with the increased prevalence of chronic pain in females.

**Fig. 6.**
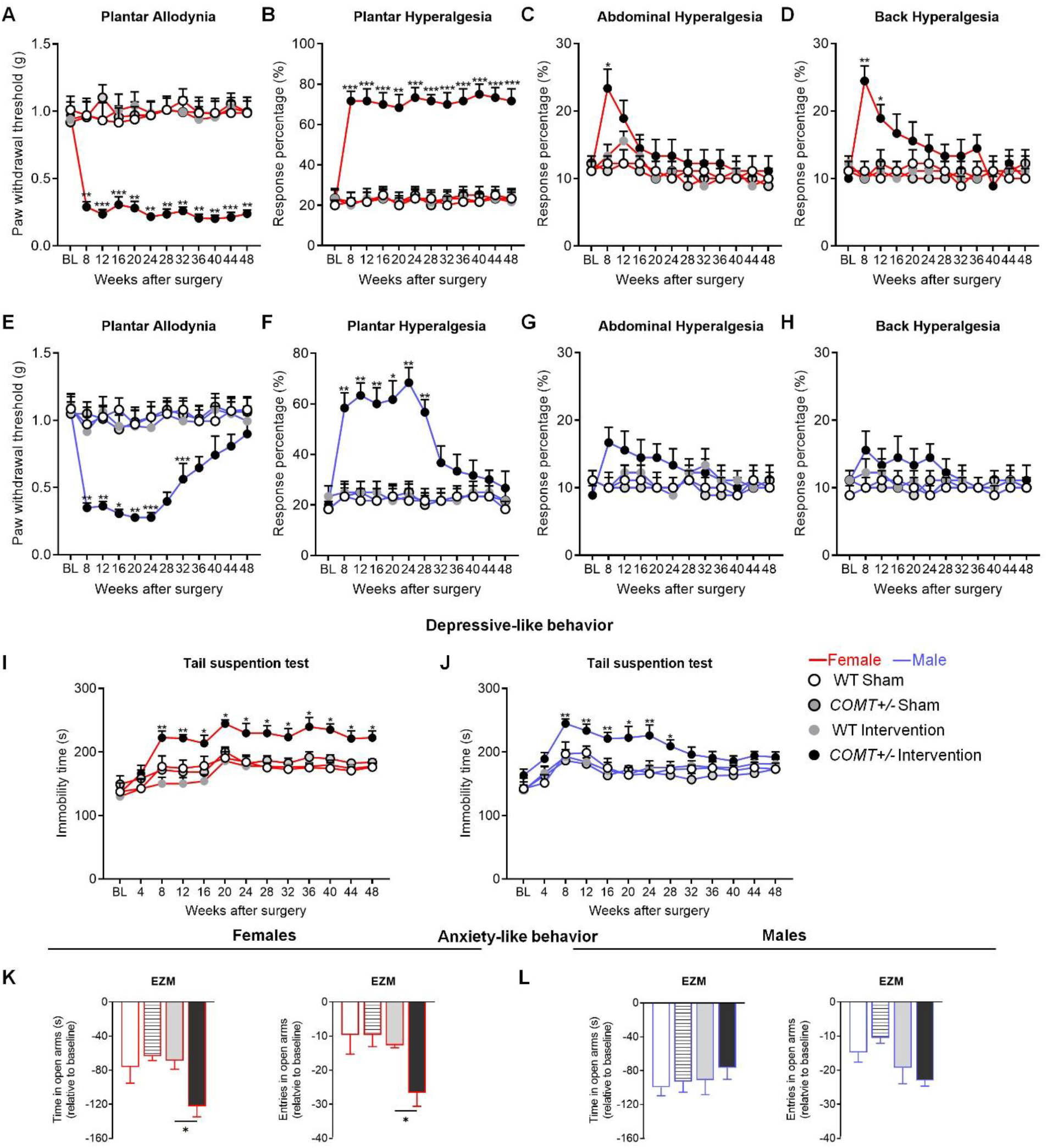
CPPC mice exhibit chronic pain and comorbid depressive and anxiety behaviors, especially females. **(A-D)** Female CPPC mice exhibit chronic mechanical allodynia and hyperalgesia at plantar sites lasting at least 48 weeks and hyperalgesia at abdominal and back sites lasting 12 weeks. **(E-H)** Male CPPC mice exhibit chronic mechanical allodynia and hyperalgesia at plantar sites, but of shorter duration (resolving by 32-36 weeks) as well as a trend towards hyperalgesia at abdominal sites at 8-12 weeks, though not significant. Both **(I)** female and **(J)** male CPPC mice exhibit comorbid depressive behavior beginning at 8 weeks that lasts at least 48 weeks for females, but resolves by 32 weeks for males. **(K)** Female, but not **(L)** male CPPC mice also exhibit anxiety-like behavior at 44 weeks. *N* = 6 mice per group. Data represent mean ± SEM. ^**^*P <* 0.01, ^***^*P* < 0.001, ^****^*P* < 0.0001 versus Sham Intervention.

Depression and anxiety are common among patients with CPPCs (*66*). We evaluated depressive-like behavior with the tail suspension test over the course of 48 weeks and anxiety-like behavior with the elevated zero maze at 44 weeks. We found that both female and male CPPC mice began to exhibit depressive behavior, evidenced by increased immobility time at 8 weeks following the intervention (long after they exhibited increased pain sensitivity). In line with their pain time course, the female CPPC mice continued to show depressive behavior at 48 weeks, while the male CPPC mice exhibited depressive behavior for 28 weeks **(Fig. 6I,J)**. Further, the female but not male CPPC mice exhibited increased anxiety levels, evidenced by the increased time and entries in open arms of the zero maze **(Fig. 6K,L)**.

### COMT genotype and stress interact to predict pain and IL-6 and IL-17A cytokine levels in a human CPPC cohort

To confirm the interaction between stress and COMT genotype in a human cohort, we examined data collected in the Complex Persistent Pain Condition (CPPC) study (*67*). This study enrolled as cases 549 individuals with one or more of the following CPPCs: episodic migraine, fibromyalgia, irritable bowel syndrome, temporomandibular disorder, and vulvar vestibulitis. Controls included 258 individuals with none of the listed pain conditions. Stress was measured using the 10 item Perceived Stress Scale (PSS) (*68*). Genotypes for the rs4633 minor T allele (in complete LD with the rs4680 ‘low activity’ minor A allele, but with superior 100% call rate) were obtained using a candidate gene panel (*69*). A total of 482 cases and 208 controls with complete data were used in the analyses.

To test the interaction between stress and the functional COMT genotype, we fit a nominal logistic model for the odds of being a CPPC case, including terms for the number of copies of the rs4633 T allele, PSS score, and a PSS x COMT genotype interaction term. The main effect of genotype was not significant (P = 0.12), but PSS was associated with an increase in risk of having a CPPC (P < 0.0001), and the interaction term was significant (P < 0.03) indicating that genotype modifies the effect of stress on the development of CPPCs (**Fig. 7A**). We then measured levels of IL-6 and IL-17A in plasma samples and found that individuals with 2 copies of the minor T allele in the high stress group had trending increases in IL-6 and significant increases in IL-17A. (**Fig. 7B,C**). This is in line with the finding from the CPPC mice and suggests that IL-6 and IL-17A may represent biomarkers for CPPCs across species.

**Fig. 7.**
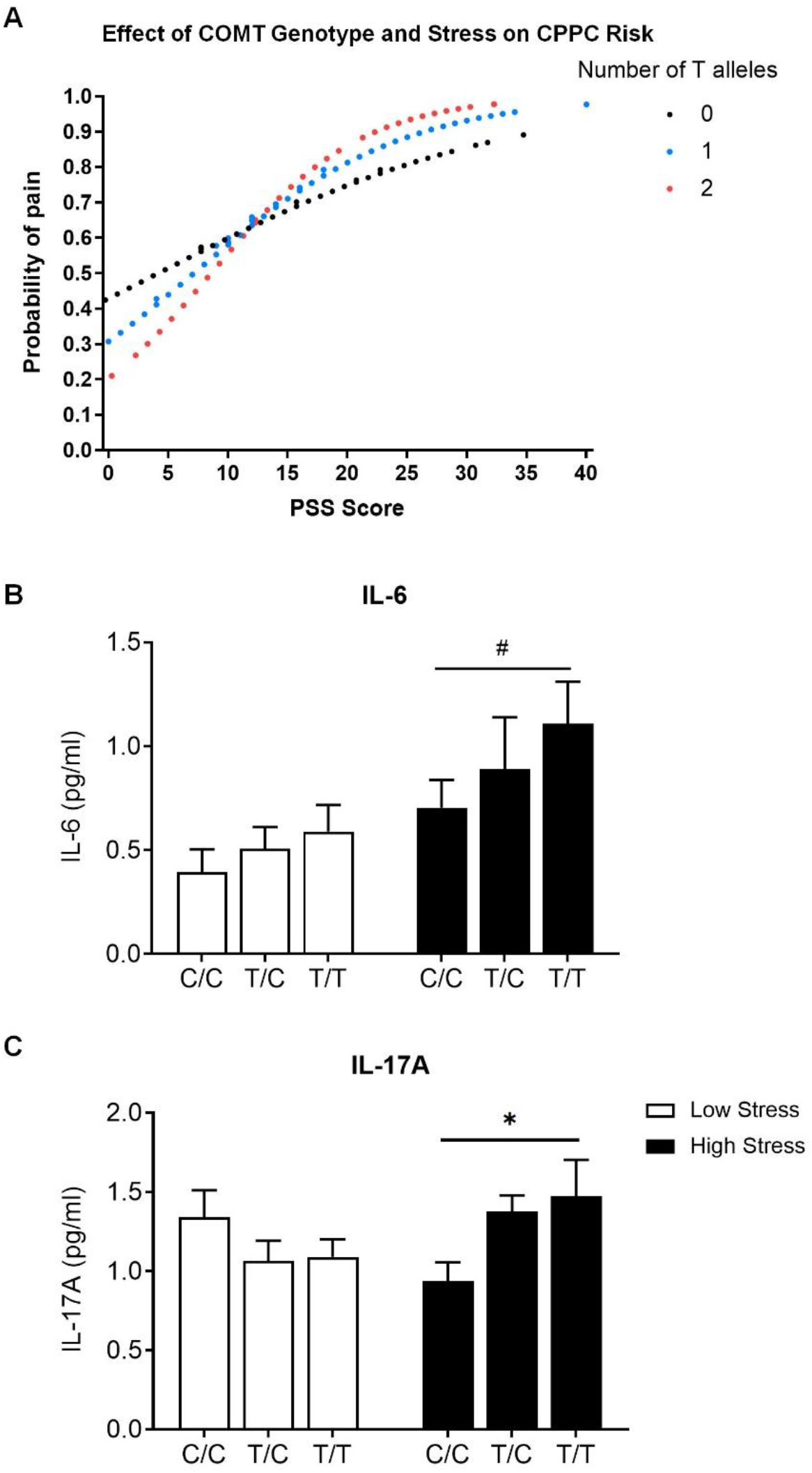
COMT genotype and stress interaction predicts CPPC risk and IL-6 and IL-17A levels in patients with CPPCs. **(A)** Data shown are predicted probabilities of pain derived from a logistic regression model for each individual in the cohort. Among individuals with higher PSS scores, CPPC risk is dose-dependently increased by the number of copies of the minor ‘low COMT activity’ T allele. Further, individuals with 2 copies of the T allele that report high stress have trending increases in plasma **(B)** IL-6 and **(C)** IL-17A levels. Data represent mean ± SEM. *P < 0.05, ^#^P < 0.1.

## DISCUSSION

CPPCs are highly prevalent conditions that tend to co-occur and have shared etiologic pathways regulated by genetic and environmental events. Yet, current animal models of CPPCs typically induce pain using chemical irritants (e.g., mustard oil, formalin, and turpentine) (*70-73*) and infectious agents (e.g., zymosan and mycobacterium) (*74, 75*) that are not implicated in the biological basis of disease. Further, current models typically focus on site-specific pain with inducers applied to the face, head, abdomen or vulvar vaginal sites and the pain measured locally (*70, 72, 74-77*). Thus, the goal of this study was to develop and validate a novel mouse model of CPPCs that incorporates clinically-relevant genetic and environmental factors to induce pain alongside an integrated approach to assess evoked pain and nociceptor activity at multiple body sites, comorbid depression and anxiety phenotypes, and cytokine biomarkers.

To induce pain, genetically predisposed COMT+/- mice were exposed to environmental stress and injury factors that, together, lead to enhanced catecholamine tone. Specifically, we found that COMT+/-, but not WT, mice exposed to a 3 day repeated swim stress paradigm followed by molar extraction surgery exhibited increased levels of NE on day 7 that persisted for at least 28 days. This finding provides evidence of physiological similarity between our model and the human CPPC disorder (*6, 7, 9*). Compared to WT mice, COMT+/- mice undergoing the repeated swim stress and molar extraction surgery intervention also exhibited pronounced multi-site body pain and depressive-like behavior lasting more than 3 months. Interestingly, while the pain was observed within just a few days following the intervention, the depressive behavior was not observed until 8 weeks later, suggesting that depression may be a consequence of pain. This finding is in line with that from a GWAS study of chronic multisite pain in 380,000 UK Biobank participants (*78*). Of 22 traits, investigators found major depressive disorder to be the most significantly correlated with pain and, using a mendelian randomization analysis, multi-site pain exposure was determined to cause depression outcome.

The COMT+/- mice undergoing the intervention also exhibited enhanced activity of primary afferent DRG nociceptors innervating hindpaw and back sites, indicative of widespread peripheral sensitization that may play a role in the chronification of pain. In addition to nociceptors, a large body of work implicates immune cells and inflammatory cytokines in the pathophysiology of CPPCs. Low COMT (*30*), stress (*31*), and injury (*32*) can produce pain by increasing the production of pro-inflammatory cytokines that sensitize nociceptors (*33-36*). Here, we identified two cytokines that were differentially expressed only in CPPC mice: IL-6 and IL-17A. IL-6 is a pro-inflammatory cytokine secreted by adipocytes, macrophages, and other cell types that stimulates inflammatory and auto-immune processes in pain, depression, and many other diseases. Circulating levels of IL-6 have been shown to be elevated in patients with CPPCs such as TTH (*44*), IBS (*45*), and FMS (*53, 79, 80*) and positively correlate with the severity of pain symptoms. Though less well characterized in the context of CPPCs, circulating levels of IL-17A have also been shown to be elevated patients with IBS (*81*) and FMS (*82*) and correlate with symptom severity. IL-17 is a pro-inflammatory cytokine secreted by activated T cells and acts in concert with other cytokines and chemokines to promote inflammation. Notably, IL-17A has been shown to positively regulate IL-6 and both IL-6 and IL-17A directly activate nociceptors and induce peripheral sensitization (*83-87*).

Beta-blockers that normalize catecholamine production (*88*) and reduce proinflammatory cytokine levels (*89*) have been shown to alleviate pain in patients with CPPCs. Previously, we found that delivery of the β3AR antagonist SR59230A prevents the development of COMT inhibitor-induced pain at plantar sites (*90*). Here, using our new model, we found that SR59230A could prevent the development of multi-site body pain and activity of nociceptors innervating the hindpaw and back in a dose dependent manner. These results demonstrate the predictive validity of our model, and highlight the importance of β3AR as a target for CPPC treatment.

In line with the female predominance of CPPCs, we found that the pain and depressive-like behavior were of greater magnitude and longer duration (lasting at least 12 months) in females compared to males. Further, only females exhibited anxiety and increased levels of IL-6. Female-specific increases in IL-6 levels have been observed in mouse models of TMD and stress (*91, 92*), suggesting it may represent a marker of sexual dimorphism. In previous work, we also found that the joint effects of COMT inhibition and stress on sexual dimorphism in pain and depressive behavior were linked to female-specific changes in the expression of brain derived neurotrophic factor (BDNF) in the spinal cord and hippocampus (*93*). BDNF is an important factor involved in nociception and depression and may also contribute to sexual dimorphism of these phenotypes.

Finally, we tested the effects of the COMT by stress interaction on pain and cytokine levels in a human CPPC case control cohort. As expected, individuals with 2 copies of the ‘low COMT activity’ T allele that reported high stress were at greatest risk for CPPCs. Intriguingly, these individuals also had increased levels of circulating IL-6 and IL-17A. To our knowledge, this is the first demonstration that levels IL-6 and IL-17A are influenced by interactions between genetic and environmental factors linked to enhanced catecholamine signaling. The fact that our novel mouse model reliably recapitulates biologically- as well as clinically-relevant features of CPPCs is a testament to its construct validity.

In conclusion, we developed and validated a novel mouse model of CPPCs that combines genetic and environmental features associated with the pathophysiologic basis of these maladaptive conditions. Based on our published and present work, we propose that interactions between low COMT activity genotype and environmental stress/injury lead to increased levels of catecholamines, which then bind to β3ARs to stimulate the production of IL-6 and IL-17A that sensitize nociceptors and ultimately lead to multi-site body pain and comorbid depression and anxiety, especially in females (**Fig. 8**). Future studies implementing this novel CPPC model are needed to directly test these and other proposed mechanisms, explore sexual dimorphism, and discover new therapeutics.

**Fig. 8.**
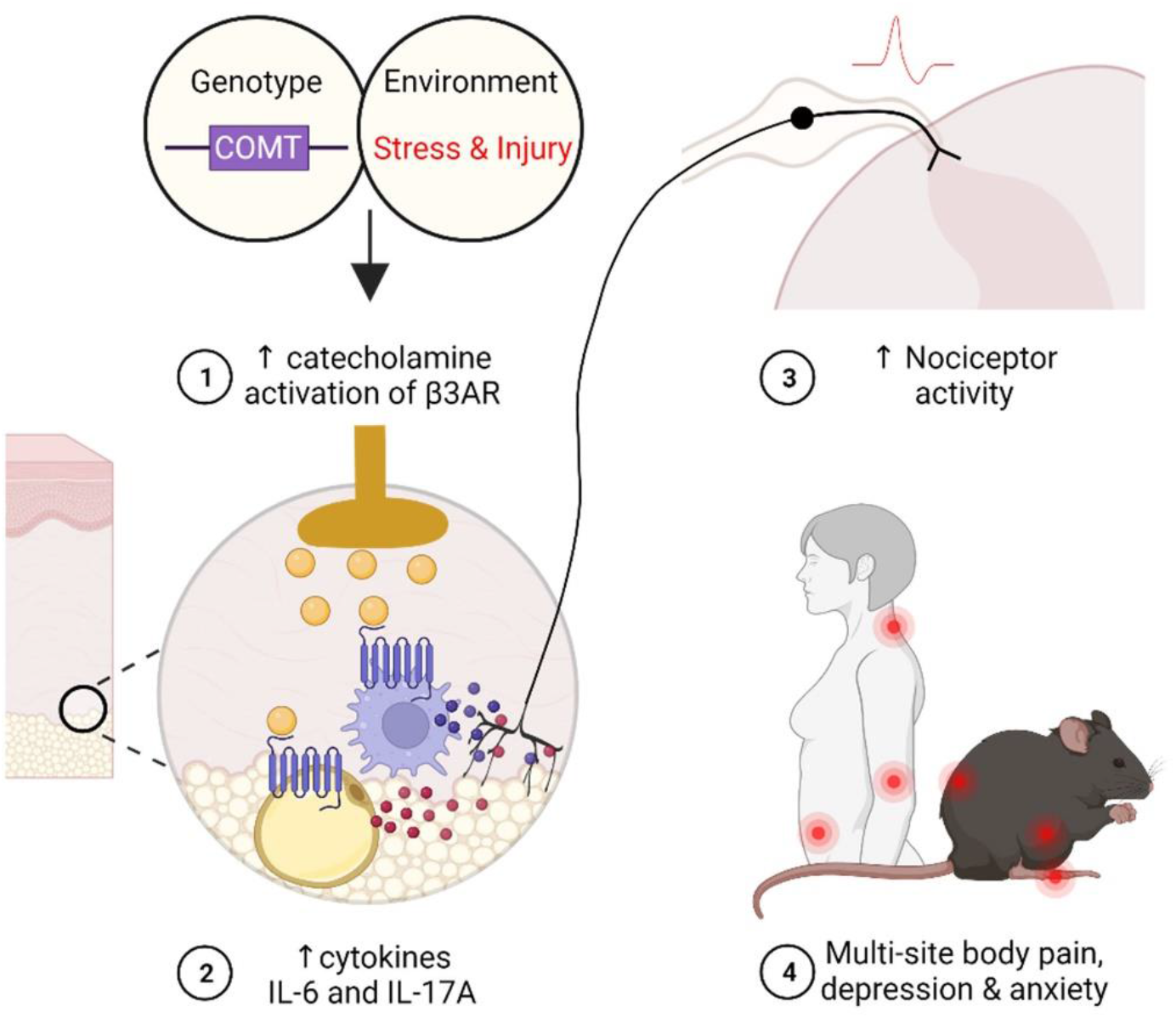
Schematic overview of proposed shared mechanisms of CPPCs in mouse and human. Interactions between low COMT activity genotype and environmental stress/injury lead to increased levels of catecholamines, which then bind to β3ARs to stimulate the production of pro-inflammatory cytokines that sensitize nociceptors and ultimately lead to multi-site body pain and comorbid depression and anxiety.

## MATERIALS AND METHODS

### Mice

Adult mice of both sexes were used for all studies. Wild-type C57BL/6J mice (Stock No: 000664 purchased from the Jackson Laboratory) and *COMT+/-* mice on a C57BL/6 background (provided by Dr. Joseph Gogos) were bred at Duke University animal facilities. Pirt-GCaMP3 mice, expressing the calcium indicator GCaMP3 in > 96% of sensory DRG neurons, were provided by Dr. Xinzhong Dong. For *in vivo* Ca^2+^ imaging studies, Pirt-GCaMP3: *COMT+/-* mice were generated by crossing Pirt-GCaMP3 mice with *COMT+/-* mice. All mice were maintained at Duke University’s animal facility and housed 2-5 per cage under a 12-hour light/dark cycle (lights on at 7 am) with food and water available *ad libitum*. All procedures were approved by the Duke University Institutional Animal Care and Use Committee (IACUC) and conformed to the National Institutes of Health Guide for the Care and Use of Laboratory Animals (*94*) and the Animal Research: Reporting of In Vivo Experiments guidelines (*95*).

### CPPC model

The swim stress model was performed as previous described (*96*). In brief, mice were placed in a glass container with 20 cm of 22-24°C water for 10 min on day 1 and 20 min on days 2 and 3. For the sham swim stress, mice underwent the same procedure with only 2 cm of water in the container. Wet mice were wiped dry and placed in a drying cage with a heat lamp above and heat pad underneath. One hour after swim stress on day 3, mice were briefly anesthetized under isoflurane, the bilateral bottom molars were removed quickly using a fin-tip needle holder. For the sham surgery, mice underwent isoflurane anesthesia and jaw/mouth opening, in the absence of molar extraction.

### Drug administration

The β3AR-selective antagonist SR59230A (2 mg/kg/day or 10 mg/kg/day; Tocris Bioscience) or vehicle (2:3 ratio of DMSO:0.9% saline) was delivered over 14 days at a constant rate by Alzet osmotic mini-pumps (model 2002; Durect Corporation, Cupertino, CA, USA). SR59230A or vehicle was administered 3 days following handling and habituation and 3 days prior to the swim stress and molar extraction intervention.

### Mechanical pain sensitivity testing

Mice were handled and habituated for 3 days prior to establishing baseline responses. On habituation and test days, mice were placed in stainless steel mesh chambers on top of a wire mesh table that permitted access to multiple body sites. **Plantar sites**. Mechanical allodynia was assessed by applying von Frey filaments (0.005-1.49 g, Stoelting) to the plantar surface of the hindpaw, using the up-down method to calculate 50% withdrawal threshold (*97-99*). Mechanical hyperalgesia was determined by measuring response frequency to a supra-threshold 0.4 g filament applied 10 times (*98*). **Abdominal and back sites**. Animals were shaved 1-2 days before the pinprick test. For the back, a mark was placed on the skin, 1*1cm^2^ above the posterior superior iliac spine. For the abdomen, a mark was placed on the skin 1*1cm^2^ surrounding the belly button. The inner needle of a spinal needle (25GX3.00N, BD) was applied to each site, and the number and type of responses to 15 stimulations recorded (*100*). Response categories include: normal/innocuous (simple withdrawal), noxious (elevating the area for extended periods of time, flinching, and licking), and null responses (no withdrawal).

### Depressive- and anxiety-like behavioral testing

Depressive-like behavior was assessed using the tail suspension test (*101*). Mice were suspended for 6 minutes from their tails, taped to a table 20–25 cm from the floor, such that they could not escape or hold onto nearby surfaces. Plastic tubes were placed over the tails to ensure mice could neither climb nor hang on their tail. Immobility time, which serves as an indicator of despair/depression, was video-recorded, and quantified by an experimenter blinded to the testing conditions. Anxiety-like behavior was assessed using the elevated zero maze. Mouse was placed on the middle of the open arm, a 5-minnute video was recorded, time and entries in the open arms were analyzed by ANY-Maze software.

### In vivo Ca^2+^ imaging

Mice were anesthetized using ketamine, xylazine and acepromazine, laminectomized to expose the lumbar region (L4-5) of the spinal cord and placed into a spinal clamp to restrict movement. After surgery and proper animal placement, sustained isoflurane inhalation was delivered such that the animal responds to hindpaw-pinch. A 10×0.5 NA objective was then focused on the DRG. Using a Zeiss 780 upright confocal microscope, a 488-nm laser line was used to image Pirt-GCaMP3-expressing nociceptors in DRG, and a 505-555 nm dichroic mirror was used to detect the emission of fluorescent signal. Following imaging controls, mice received noxious mechanical stimulation (200-300g) on L5 DRG innervated hind paw by Pincher Analgesia Meter (IITC, 2450), or a spinal needle poke on L4 DRG innervated back skin. Live images were acquired at ∼8-10 frames in frame-scan mode per 8–10 s. Images were acquired by a researcher blinded to experimental condition and processed with Fiji to compare and count the number of evoked neurons pre- and post-noxious stimuli.

### Norepinephrine and cytokine measurements

To detect plasma NE, the whole blood was collected in the cardiac puncture after endpoint or baseline with EDTA coated tube. NE ELISA kit was purchased from the Eagle Biosciences (Cat# NOR31-K01). ELISA assays were conducted according to the manufacturer’s instructions. The absorbance was measured in 450 nm wavelength using a 96 well xMark™ Microplate plate reader (BioRad). The cytokine levels were calculated by standard curve and analyzed using microplate manager 6 software (BioRad). Cytokines were measured with MSD plates and performed according to the manufacturer’s instructions.

### CPPC cohort

The human CPPC study enrolled 549 individuals with 1 or more CPPC: episodic migraine, temporomandibular disorder, irritable bowel syndrome, vulvodynia, and fibromyalgia. Controls included 258 individuals with none of the listed pain conditions. Stress was measured using the 10-item perceived stress scale (PSS) (*68*). Genotypes for 10 SNPs in the COMT locus were obtained using a candidate gene panel. The call rate of the Val158Met variant (rs4680) was only 70%, but a nearby SNP in high LD (rs4633, R2 = 1.0 in CEU) with rs4680 had a call rate of 100% and was used as a proxy for Val158Met; the T allele of rs4633 is functionally equivalent to the Met allele of rs4680 characterized by lower enzymatic activity. A total of 482 cases and 208 controls with complete data were used the COMT x stress interaction analyses (**Table S1**). Samples from a subset of 40 CPPC cases and 40 controls were used in the cytokine analyses.

### Statistical analyses

Statistical analyses of animal data were performed using GraphPad Prism 9 (GraphPad Software, La Jolla, CA, USA). Group differences in mouse behavioral responses, evoked cell numbers in *in vivo* Ca2+ imaging, and cytokine levels were analyzed by 1-way or 2-way ANOVA. Post hoc comparisons were performed using the Bonferroni test, which is corrected for multiple comparisons. Statistical significance was defined as *P* < 0.05. Statistical analyses of human data were performed using SAS 9.4 (SAS Institute Inc., Cary, NC, USA). To test for COMT genotype and stress interactions in the CPPC cohort, we fit a nominal logistic model for the odds of having a CPPC, including terms for the number of copies of the rs4633 T allele, PSS score, and a PSS x COMT genotype interaction term.

## List of Supplementary Materials

Fig. S1. Establishment of chronic primary pain conditions model in COMT+/- mice

Fig. S2. β3AR antagonist prevented the mechanical pain sensitivity in COMT+/- CPPC mice

Fig. S3. CPPC model in COMT+/- mice exhibits chronic pain phenotype

## Acknowledgments

We would like to thank all of the mouse and human study participants for their invaluable contributions to this work. We also thank the late William Maixner for his vision, stewardship, and tireless support of our translational CPPC research efforts.

## Author contributions

Conceptualization: AGN, XZ, YW

Mouse study methodology and investigation: YW, XZ, SHK, MK, JC, EG

Human study methodology and investigation: GDS, AB, and SS

Result interpretation: YW, XZ, AB, and AGN

Writing- original draft: YW and AGN

Writing- review and editing: YW, XZ, SHK, MK, JC, EG, SS, AB, AGN

## Data and materials availability

All data associated with this study are present in the paper or the Supplementary Materials.

