## Supplemental Fig. 1 ∼ Fig. 3 for "A Novel Mouse Model of Chronic Primary Pain Conditions that Integrates Clinically Relevant Genetic and Environmental Factors"

**Supplementary Materials**

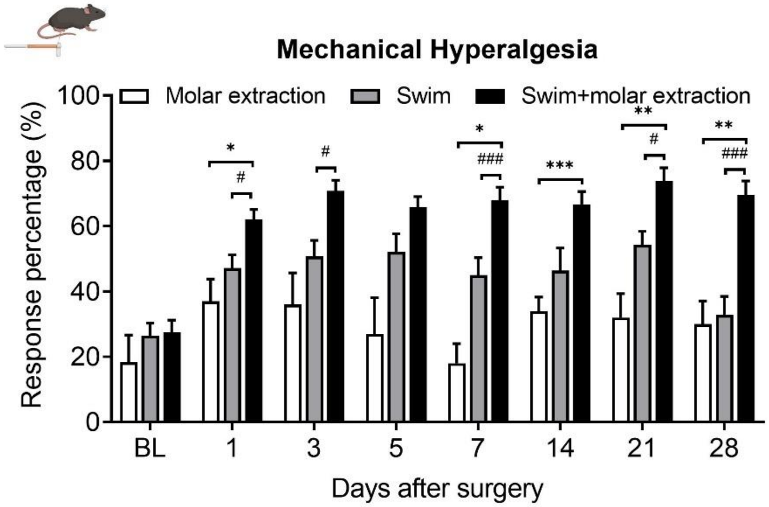


**Fig. S1. Swim stress and injury exhibit robust and sustained mechanical hypersensitivity at plantar sites.** Mice undergoing swim and molar extraction exabit mechanical hyperalgesia compared to mice undergoing swim stress or molar extraction alone. *N* = 5-12 mice per group. Data represent mean ± SEM. $^{*}P < 0.05$, $^{**}P < 0.01$, $^{***}P < 0.001$ versus Molar extraction. $^{\#}P < 0.05$, $^{\#\#\#}P < 0.001$ versus Swim.

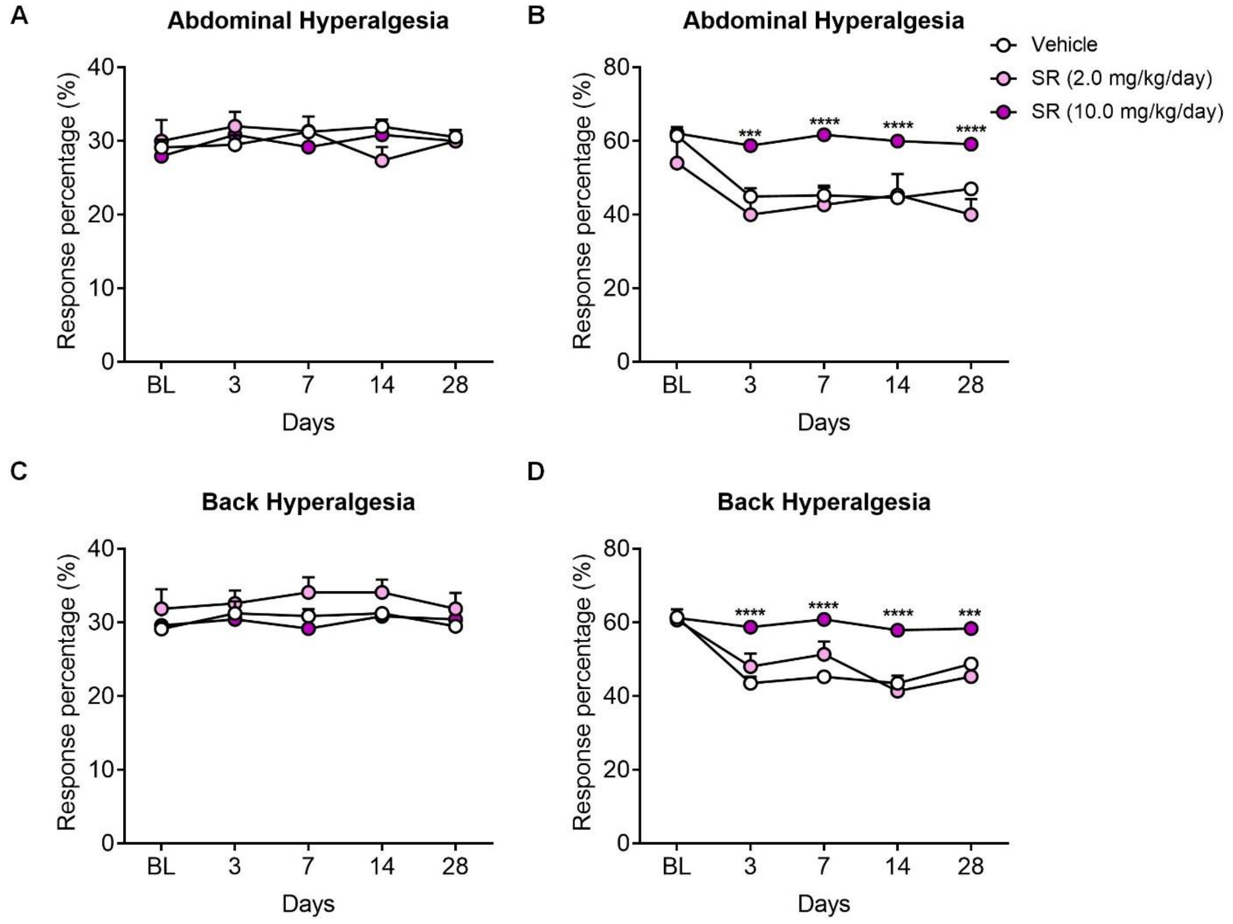


**Fig. S2. The β3AR antagonist SR59230A reduced multi-site mechanical hypersensitivity at abdominal and back sites.** Compared to naïve mice, SR59230A dose-dependently reversed mechanical hypersensitivity at abdominal evidenced by unchanged innocuous responses (**A**) and increased null responses (**B**) at abdominal sites, and unchanged innocuous responses (**C**) and increased null responses (**D**) at back sites among female and male CPPC mice. N = 19 (11M, 8F) in Vehicle, N=10 (5M, 5F) in SR (2.0 mg/kg/day), and N = 16 (7M, 9F) in SR (10.0 mg/kg/day) group. Data represent mean ± SEM. $^{***}P < 0.001$, $^{****}P < 0.0001$ versus vehicle or naïve.

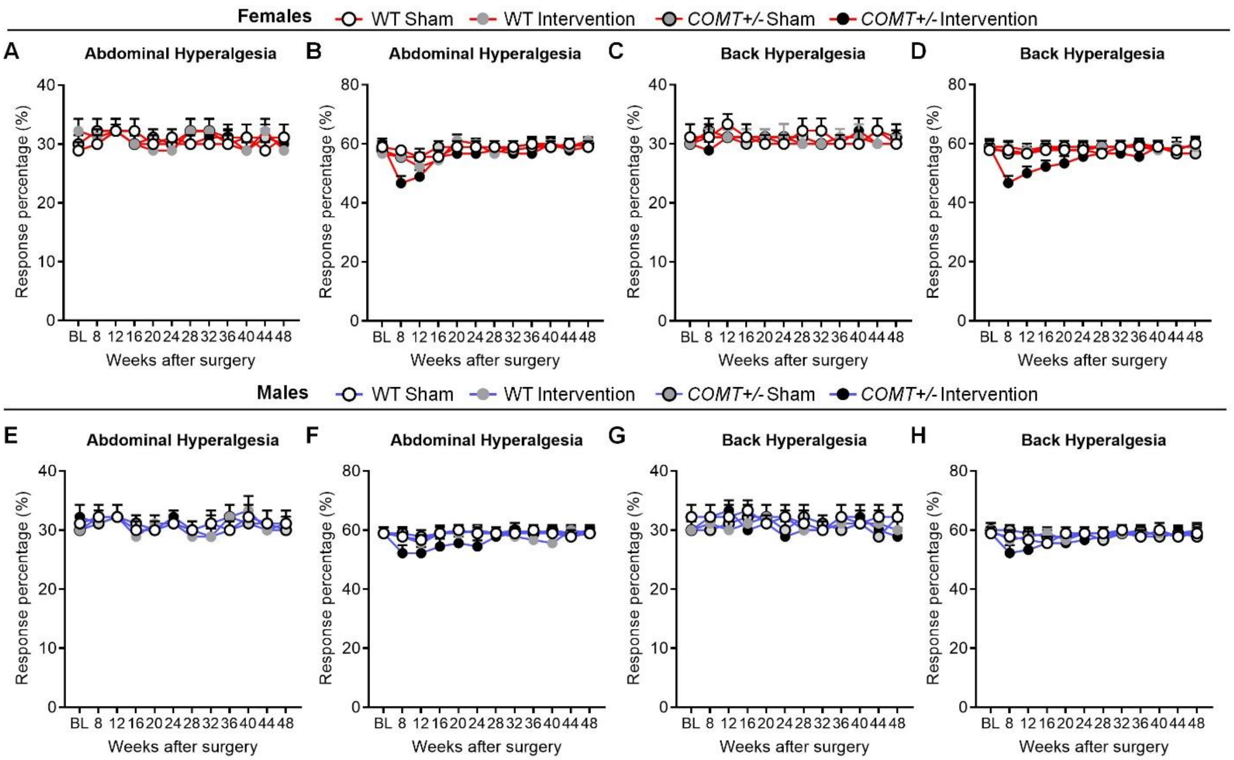


**Fig. S3. CPPC mice exhibit chronic pain behaviors.** The innocuous response (A) and null responses (B) at abdominal sites and innocuous response (C) and null responses (D) at back sites in Female CPPC mice remain unchanged. The innocuous response (E) and null responses (F) at abdominal sites and innocuous response (G) and null responses (H) at back sites in male CPPC mice remain unchanged at 8-12 weeks. *N* = 6 mice per group. Data represent mean ± SEM.
